## Supplementary Figure 1 and Table 1 for "Ventromedial prefrontal cortex compression during concept learning"

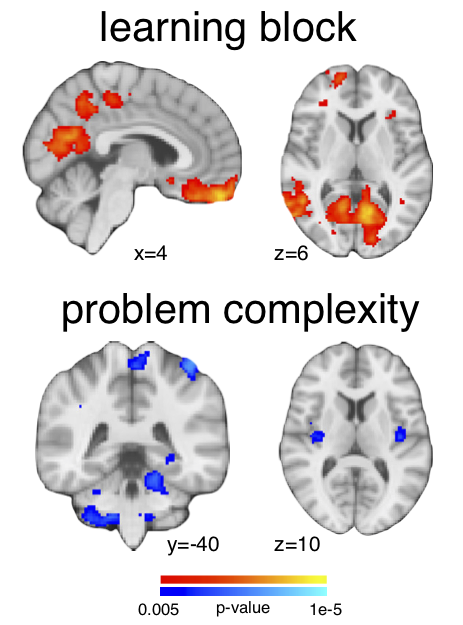

**Supplementary Figure 1:** Whole brain maps of neural compression analysis for main effects of learning block and problem complexity (N=23). Neural compression increased with learning block (top) in several regions including vmPFC, posterior cingulate, and retrosplenial cortex. Neural compression decreased with higher problem complexity (bottom) in bilateral insula, superior parietal, and cerebellum (see Supplementary Table 1 for all identified clusters). These maps were generated with a voxel-wise threshold of p = 0.005 and cluster-extent threshold of p = 0.05.

|  | peak location | | peak *t* value | size | labels |
| --- | --- | --- | --- | --- | --- |
| learning block | | 30, -78, 38 | 4.56 | 20162 | occipital, inferior parietal, angular gyrus |
|  |  | 42, 22, -42 | 4.92 | 10450 | bilateral anterior temporal, ventromedial prefrontal cortex |
|  |  | -18, 28, 42 | 3.62 | 671 | left superior frontal gyrus |
|  |  | 22, -88, -32 | 3.81 | 244 | right cerebellum |
| problem complexity | | -14, -22, -34 | 3.7 | 1462 | cerebellum, brain stem |
|  |  | 34, -14, 14 | 3.33 | 744 | right insula |
|  |  | -42, -42, 62 | 3.9 | 537 | left superior parietal |
|  |  | -2, -30, 66 | 3.48 | 524 | bilateral precentral gyrus |
|  |  | -40, -14, 6 | 3.4 | 411 | left insula |

**Supplementary Table 1:** Significant clusters showing main effects of learning block or problem complexity from the neural compression searchlight analysis. Cluster information includes the peak voxel location in MNI coordinates (x, y, z), the *t* statistic of the peak voxel, the cluster size in number of voxels, and the corresponding anatomical label(s) as defined in the Harvard-Oxford Structural Atlas.
